# Regulatory T cells limit age-associated retinal inflammation and neurodegeneration

**DOI:** 10.1101/2023.10.11.561874

**Authors:** María Llorián-Salvador, Alerie G de Fuente, Christopher E. McMurran, Rosana Penalva, Yvonne Dombrowski, Alan W. Stitt, Denise C Fitzgerald

## Abstract

Aging is the principal risk factor for retinal degenerative diseases, which are the commonest cause of blindness in the developed countries. These conditions include age-related macular degeneration or diabetic retinopathy. Regulatory T cells play a vital role in immunoregulation of the nervous system by limiting inflammation and tissue damage in health and disease. Because the retina was long-considered an immunoprivileged site, the precise contribution of regulatory T cells in retinal homeostasis and in age-related retinal diseases remains unknown. Our study shows that regulatory T cell elimination leads to retinal pigment epithelium cell dysmorphology, and accumulation of phagocytes in the subretinal space of young and aged mice. However, only aged mice experience retinal neurodegeneration and gliosis. Surprisingly, adoptive transfer of young but not aged regulatory T cells reverse these changes. This study reveals a previously unknown protective role of regulatory T cells in maintaining aged retinal homeostasis.

## Introduction

Aging represents the most significant risk factor for various sight-threatening retinal diseases, such as diabetic retinopathy, age-related macular degeneration (AMD) or glaucoma. Although the precise pathobiology underpinning age-related retinal degeneration remains ill-defined, there is broad recognition of the crucial role played by chronic low-grade chronic inflammation(Xu et al. 2009; Kaur & Singh 2021). Historically, the retina has long been regarded as immunoprivileged. This privileged status is maintained by protective retinal-blood-barriers such as the retinal pigment epithelium (RPE) and a localized immune system including microglia, which prevent the entry of pathogens and systemic inflammation into the retina(Chen et al. 2019). The aging retina is characterized by chronic low-grade inflammation (also referred to as “para-inflammation” or “inflammaging”) resulting in age-mediated endogenous insults and impaired defense mechanisms (i.e. reduced phagocytic activity, migration from microglia, disruption of blood-retinal barrier, etc). However, it is now recognised that age-related immune dysregulation contributes to retinal degeneration even in the absence of overt disease, challenging the notion of complete immune privilege in the retina. This dysregulation leads to detrimental age-related pathologies such as AMD, diabetic retinopathy or glaucomatous retinopathy(Chen et al. 2019; Chen et al. 2011).

Regulatory T cells (Treg) maintain and restore tissue homeostasis by regulating immune responses and inflammation and promoting tissue regeneration in a range of tissues(Muñoz-Rojas & Mathis 2021). In the central nervous system (CNS), Tregs have key neuroprotective functions by modulating microglial and astrocyte activation to prevent gliosis and inflammation and enhance remyelination to limit axonal degeneration(Shi et al. 2021; Yshii et al. 2022; Ito et al. 2019; Dombrowski et al. 2017). Treg density in healthy retinal parenchyma is very low, although Treg infiltration of the neuropil occurs during retinal inflammatory diseases(McPherson et al. 2012). During acute retinal inflammation, such as in uveitis, Tregs exert immunosuppressive functions which counterbalance uncontrolled immune responses and thereby limit degenerative pathology(McPherson et al. 2012). Ischemia also increases Treg density in the retina where they participate in preventing supra-retinal damage and fostering reparative intra-retinal angiogenesis(Deliyanti et al. 2017), demonstrating that Treg play important roles in retinal disease. Yet the role of Treg in age-associated retinal pathology has been largely unexplored.

As we age, levels of Treg increase in blood(Garg et al. 2014; Elyahu et al. 2019), but whether this supports immune regulation in the aged retina is unknown. To address this, we asked whether genetic ablation of Treg impairs retinal homeostasis in young and aged mice. Here, we show that Treg depletion leads to accelerated retinal neurodegeneration and gliosis in aged but not young mice. Thus, we demonstrate that Treg have a pivotal role in preventing aged retinal neurodegeneration and subsequent gliosis. We have also shown that young Treg are more efficient than aged Treg in reversing age-related retinal degeneration. This work sets the groundwork to investigate Treg immunosuppressive and neuroprotective capacity as an effective treatment for age related retinal degenerative diseases.

## Results

### Absence of aged Treg leads to retinal neurodegeneration

To determine the role of Treg in the adult and aged retina in the absence of apparent disease, we depleted Treg using diphtheria toxin (DT) in young (<4 months) and aged (>18months) B6.129(Cg)-*Foxp3*^*tm3(DTR/GFP)Ayr*^/J(FoxP3-DTR) mice(Kim et al. 2007; Dombrowski et al. 2017) (**Fig. 1A, Fig EV1A, B**) and evaluated changes in the neural retina and retinal pigment epithelium (RPE). Histological analysis at 2.5 weeks post-Treg depletion identified a significant decrease in total retinal thickness in aged but not young mice (**Fig EV1C, D**). On further analysis, no changes in photoreceptor nuclei were observed in young mice lacking Treg, while aged mice had a significant reduction of total photoreceptors (DAPI^+^ rows in the ONL) following Treg depletion (**Fig. 1B, C**). Specifically, photoreceptor loss was associated with a decrease in both, Cone arrestin^+^ (CA^+^)cones (**Fig. 1B, D**) and CA^-^ rods (**Fig. 1B, E**). To further investigate Treg depletion-associated neurodegeneration, we examined the second-order neurons in the retina. In normal retinas, the bodies of rod bipolar cells reside in the outer region of the inner nuclear layer (INL). These cells exhibit a cluster of dendrites that extended into the outer plexiform layer (OPL). Treg depletion did not affect overall rod or cone bipolar cell number in young or aged Treg-depleted mice (**Fig. 1F-J**). However, in PKCα^+^ rod-bipolar cells and secretagogin^+^ cone-bipolar cells, Treg depletion in aged mice altered cell body laminarity (**Fig. 1F, I**). This was more pronounced in PKCα^+^ rod-bipolar cells; in which despite unaltered total number of somas, there was a significant increase of ectopic PKCα^+^ cell bodies located below the outer plexiform layer (OPL) due to shorter axons (**Fig. 1F, G**). Upon Treg depletion, we also found bipolar dendrites and synaptophysin sprouts expanding from the OPL towards the ONL (**Fig. 1F**). This cellular remodelling (consisting of the retraction of the bipolar cells, sprouting of dendrites and formation of ectopic synapses) has been previously observed in diseases associated with photoreceptor loss(Cuenca et al. 2014). We did not observe any alteration in retinal ganglion cells across experimental groups (**Fig EV1E-G**) suggesting that neurodegeneration following Treg depletion in aged mice predominantly impacted the outermost layers of the retina.

**Figure 1:**
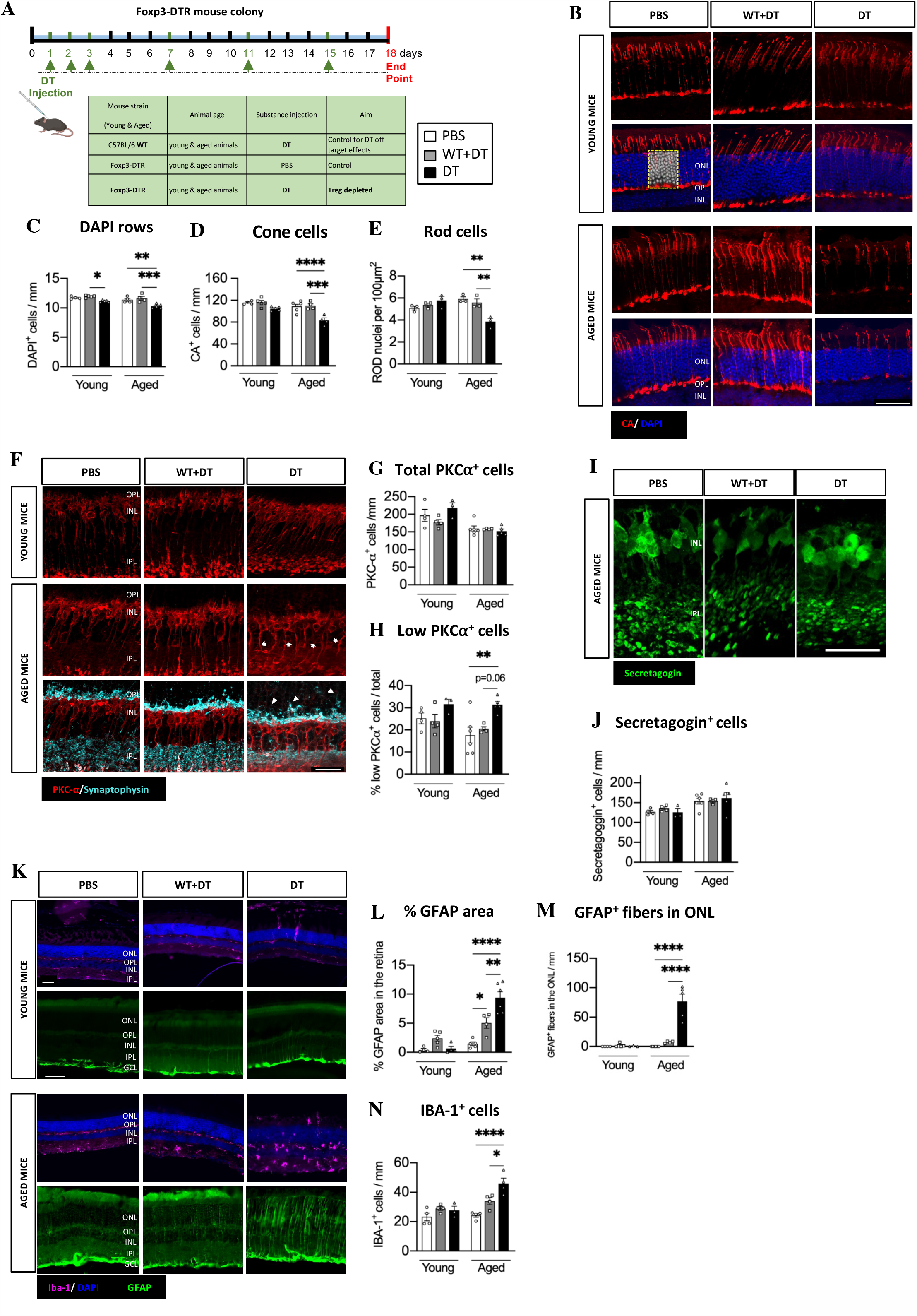
Treg depletion in aged but not young mice accelerates retinal neurodegeneration. **A** Diagram of the treatment regime and research groups. **B-E** Immunostaining and quantitative analysis of photoreceptors. Representative images showing photoreceptors in the ONL (**B**) (Cones, CA^+^; Rods DAPI^+^ CA^-^) in young and aged Foxp3-DTR (scale bar= 50 μm). Quantitative analysis of the number of rows of DAPI^+^ nuclei in the ONL (C). Quantification of the number of cone photoreceptor cells (**D**) (CA^+^ cells). The number of rods (**E**), determined by deduction of CA^+^ cells from the total number of cells (DAPI^+^ cells in ONL). **F-J** Immunostaining and analysis of rod bipolar cells (**F-H**) and cone bipolar cells (**I, J**). Representative image of PKC-α (**F**) (red) and synaptophysin (blue, lower panels). Arrows indicate abnormal location of some PKC-α^+^ cell soma lower within the INL. Arrowheads indicate abnormal rod bipolar dendrite and synaptic vesicles sprouts into the ONL (scale bar=25 μm). Quantitative analysis of the number of PKC-α^+^ cells (**G**) (n=3-6 mice, mean±s.e.m shown, 1-way ANOVA followed by Bonferroni’s multiple comparisons test) and (**H**), percentage of PKC-α^+^ somas found in a lower ectopic location out of the total of PKC-α^+^ cells (n=3-6 mice, mean±s.e.m shown, 1-way ANOVA followed by Bonferroni’s multiple comparisons test). Representative image (**I**) (scale bar= 25 μm) and quantification of rod bipolar cells in the INL (**J**) using secretagogin staining. **K-N** Analysis of IBA-1^+^ microglial cells and GFAP^+^ Muller cells determined by immunohistochemistry. Representative image of microglial cells (**K**) (IBA-1^+^) and gliotic Muller cells (GFAP^+^ fibers) (scale bars=50 μm). Quantitative analysis of the percentage of total GFAP^+^ area (**L**), the number of fibers GFAP^+^ in the ONL (M) and total number of IBA-1^+^ cells in all layers of the retina (**N**). Data information: B-N, n=3-6 mice, data presented as mean±s.e.m. ^*^P<0.05; ^**^ P<0.01; ^***^P<0.005; ^****^p<0.001, 1-way ANOVA followed by Bonferroni’s multiple comparisons test.

### Retinal gliosis is associated with the lack of Treg in the aged retina

To determine if Treg depletion affected retinal glia, we first examined Müller glial cell responses. In the absence of Treg we observed a significant increase in GFAP^+^ staining in the neuroretina (Fig. 1K) in aged but not young Treg-depleted mice, suggesting an intense Müller cell gliosis. In addition to overall increased GFAP expression (GFAP^+^ area of retina) (**Fig. 1K, L**) we also observed an expansion in reactive Müller cell number in the ONL (**Fig. 1K, M**). Müller cell hypertrophied branches could be a compensatory response growing towards the outer limiting membrane to compensate for photoreceptor loss in an attempt to maintain retinal cytoarchitecture, as described in other retinal diseases characterised by photoreceptor loss, such as retinitis pigmentosa. Hyperreactivity has also been observed in other retinal diseases characterised by photoreceptor loss such as retinitis pigmentosa(Fernández-Sánchez et al. 2015).

We next examined whether Treg depletion modified microglial activity in the neuroretina. Microglia are the localized immune cells in the retina, and they show an increased activation in the aged retina(Xu et al. 2009; Chen et al. 2019). While Treg depletion in young mice did not affect retinal microglia, a significant increase in the density of IBA-1^+^ microglia was observed in the neuroretina of aged mice (**Fig. 1K, N**). This increase was detected in all retinal layers, but most significantly in the ONL which is normally devoid of microglia (**Fig EV2A**). IBA-1^+^ cell increase was mostly related to ameboid-shaped microglia (**Fig EV2B**), a morphological and phenotypic shift usually associated with pro-inflammatory responses(Madry et al. 2018). To validate this phenotype, we quantified the number of MHCII^+^ pro-inflammatory microglia (Frank et al. 2006; Butovsky & Weiner 2018) in aged Treg-depleted mice and observed a significant increase of these IBA-1^+^MHCII^+^ microglia in the neuroretina (Fig EV2C, D).

These results indicate that Treg are necessary to prevent photoreceptor death and distorted lamination of bipolar cells in aged retinas. This appears to be linked with Müller cell hypertrophy as well as the accumulation of pro-inflammatory microglia in the nuclear layers of the retina.

### Treg adoptive transfer prevents retinal degeneration and rescues gliosis

We next sought to determine if Treg reconstitution could rescue age-associated retinal neurodegeneration. To do so, we adoptively transferred purified young and aged Treg into Treg-depleted aged mice to determine whether aging affects the capacity of Treg to maintain retinal normal immunoregulation and limit the negative effects of an uncontrolled inflammation in the aged neuroretina (**Fig. 2A**). We first examined if reconstitution with young and aged Treg diminished photoreceptor loss. Surprisingly, transfer of young but not aged Treg rescued photoreceptor (cone and rod) loss in the neuroretina (**Fig. 2B-E**). Although no differences were detected in the number of rod and cone bipolar cells upon Treg depletion (**Fig. 2F, G, H, J**), adoptive transfer of young but not aged Treg improved the bipolar lamination pattern, showing a decrease in the number of ectopic PKCα^+^ somas, as shown by the diminished number of PKCα^+^ cell bodies with a lower nuclear location (**Fig. 2F, I**). Additionally, young Treg also succeed in reducing the bipolar dendrite sprouting into the ONL (**Fig. 2F, I**). We next evaluated if adoptive transfer of Treg in aged mice inhibited Müller cell gliosis. In line with what we observed for photoreceptor loss, only young but not aged Treg transfer prevented Müller cell gliosis in Treg-deficient aged retinas (**Fig. 2K, L, M**), supporting the concept of Müller cell gliosis being a compensatory response to retinal degeneration as described previously(Bringmann et al. 2009; Hippert et al. 2015). Lastly, we studied the effect of Treg adoptive transfer on microglia density in the neuroretina. In contrast to what we observed with Müller cells, both young and aged Treg reduced the number of total IBA-1^+^ microglia as well as the number of IBA-1^+^MHCII^+^ microglia present in the neuroretina (**Fig. 2K, N, Fig EV2A-D**). Therefore, since adoptive transfer of aged Treg reduces the number of microglia in the neuroretina, but not photoreceptor loss, these data indicate that the increased microglia in the neuroretina is not triggering photoreceptor loss and neurodegeneration at the outermost layers of the retina.

**Figure 2:**
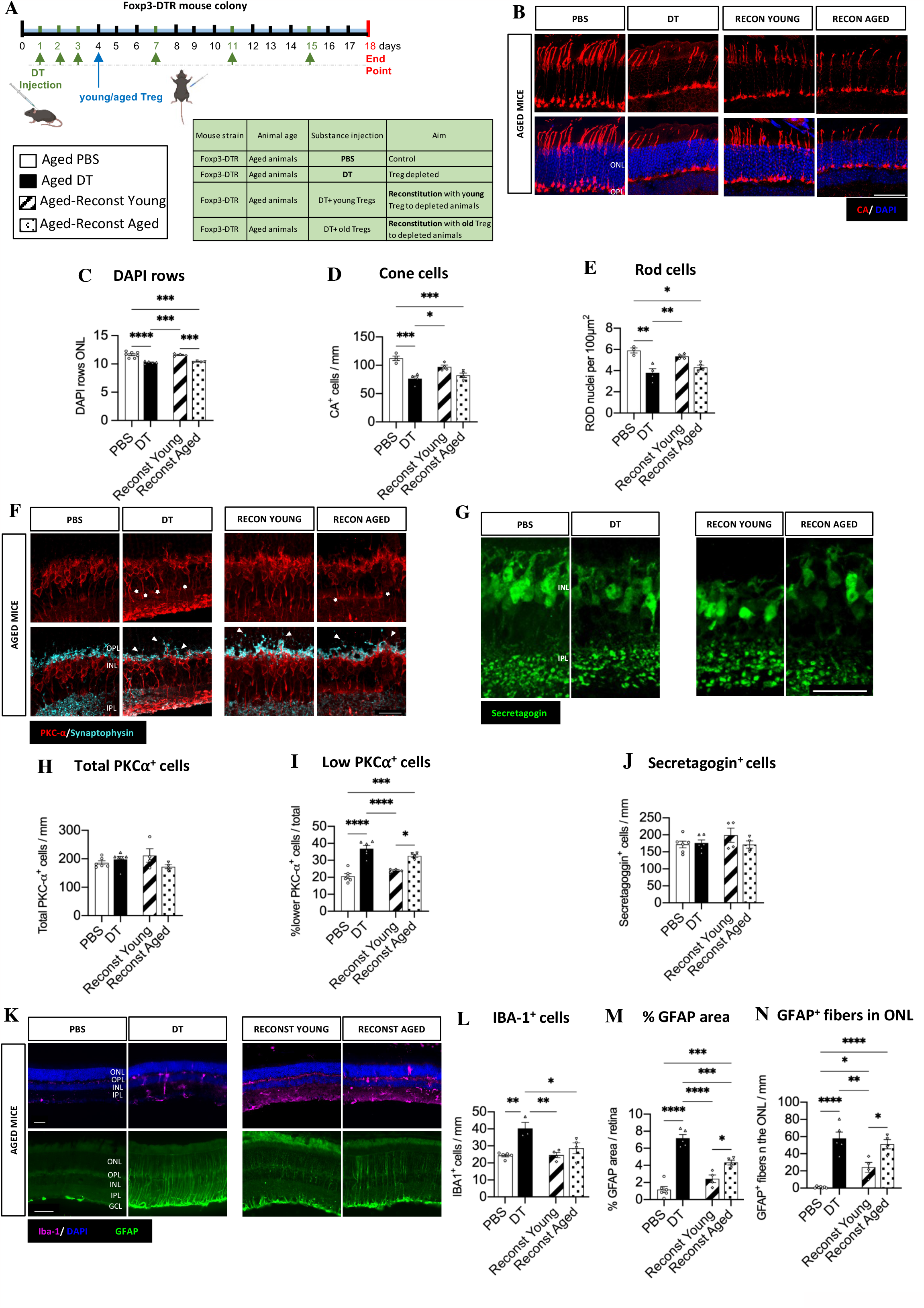
Adoptive transfer of young but not aged Treg rescues aged retinal neurodegeneration. **A** Diagram showing the experimental design. **B-E** Immunostaining and analysis of the effect of Treg adoptive transfer on photoreceptors. Representative images (**B**) (scale bar = 50 μm) and quantification photoreceptors (**C-E**). Total photoreceptors (**C**), CA^+^ cones (**D**) and CA^-^ rods (**E**) in the ONL of aged control, Treg depleted and Treg reconstituted aged retinas. **F-J** Alterations in bipolar cells after Treg adoptive transfer in aged animals. Representative images (**F**) (scale bar = 25 μm) and quantification of total rod bipolar cells (**H**) and rod bipolar cells with a shorter axon and displaced nuclei (**I**) identified by PKC-α (red) and synaptophysin (cyan) immunostaining (n=4-6 mice, mean±s.e.m shown, 1-way ANOVA followed by Bonferroni’s multiple comparisons test). Representative images (G) (scale bar=25 μm) and quantification of rod bipolar cells (J) identified by secretagogin immunostaining. **K-N** Changes in Muller cell gliosis and microglia in aged animals after the adoptive transfer of Treg. Representative image showing microglia (purple) and Müller cell (green) gliosis (**K**) (Scale bar=50 μm). Quantification of total IBA-1^+^ microglia (**I**) (n=3-5 mice, mean, 1-way ANOVA followed by Bonferroni’s multiple comparisons test). Quantification of GFAP^+^ area (**M**) and the number of GFAP^+^ fibers crossing the ONL (**N**) in the aged retina Data information: B-N, n=3-7 mice, data presented as mean±s.e.m. ^*^P<0.05; ^**^ P<0.01; ^***^P<0.005; ^****^p<0.001, 1-way ANOVA followed by Bonferroni’s multiple comparisons test.

### Treg maintain RPE integrity and limit phagocyte accumulation in the subretinal space

It was previously reported that a range of retinal pigment epithelium (RPE) defects occur in the aging retina. With age, RPE cells show an altered morphology, which includes discontinuity of the cytoskeletal bands between adjacent cells and altered cytoarchitecture (appearance of enlarged and irregular cells), contributing to altered retinal immune regulation and enhanced chronic low-grade inflammation described in the aged retina(Tarau et al. 2019). In agreement with this, we also observed significantly more enlarged RPE cells in aged animals (**Fig. 3A, B**). Such age-related RPE pathology, was exacerbated by Treg depletion and even developed in young mice lacking Treg (**Fig. 3A, B**). In addition, CD68^+^ phagocytes were increased in the subretinal space of both, young and aged mice upon Treg depletion (**Fig. 3C, D**). As per our observations in the neuroretina, phagocyte infiltration in the subretinal space mainly consisted of IBA-1^+^MHCII^+^ proinflammatory microglia/macrophages (**Fig. 3E, F-J**). Thus, Treg depletion led to accumulation of innate immune cells in the subretinal space and RPE alterations in both young and aged mice. However, these alterations were associated with neurodegeneration and gliosis only in the aged retina. Taking into account that neurodegeneration is present mostly in the outermost part of the retina, these data suggest that aged retinas are more susceptible to the neurotoxic effects of accumulated pro-inflammatory innate immune cells in the subretinal space. Hence, aged retinas are more dependent on Treg for retinal immunoregulation and maintenance of tissue homeostasis. Interestingly, transfer of young Treg diminished the RPE cell morphological changes, while transferred aged Treg failed to do so efficiently (Fig. 3A, B). Similarly, transfer of aged Treg did not limit CD68^+^ and IBA-1^+^MHCII^+^ cell infiltration in the subretinal space, while transfer of young Treg was highly efficient at doing so (Fig. 3C-F). Therefore, Treg are essential to limit age-related RPE dysmorphology and innate cell accumulatio in the subretinal space. However, in older age Treg have a limited capacity to control an already established retinal inflammatory environment.

**Figure 3:**
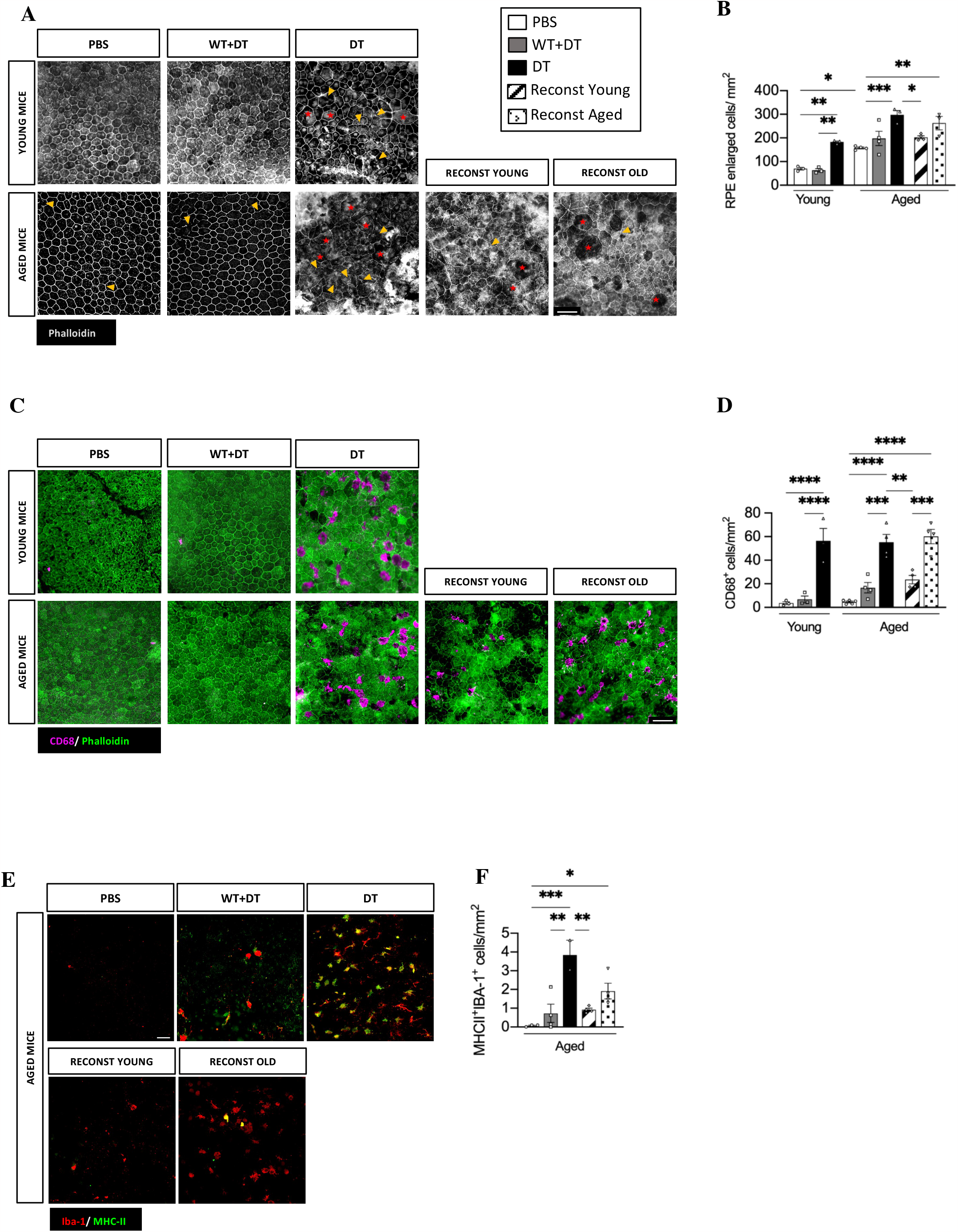
Treg depletion leads to blood-retinal barrier damage and innate immune cell infiltration in the subretinal space of young and aged mice. **A, B** Analysis of RPE cell changes. Representative image of phalloidin (**A**) indicating the geometry of RPE cells. Stars indicate enlarged cells and arrowheads fragmented cytoskeleton (scale bar=50 μm). Quantification indicating the number of RPE cells with an enlarged morphology per area of RPE/choroid flatmount (**B**). **C, D** Determination of the number of CD68^+^ cells in subretinal space. Representative image of CD68 and phalloidin immunostaining in the RPE/choroid flatmount (**C**) (scale bar=50 μm) and (**D**) quantitative analysis of the number of CD68^+^ cells in the subretinal space. **E, F** Determination of CD68^+^MHCII^+^ cells in the subretinal space. Representative image (**E**) (scale bar=50 μm) and (**F**) quantification of CD68^+^MHCII^+^ innate immune cells in the subretinal space upon Treg depletion and reconstitution in aged mice (scale bar= 50 μm). Data information: A-F, n=2-5 mice, data presented as mean±s.e.m. *P<0.05; ** P<0.01; ***P<0.005; ****p<0.001, 1-way ANOVA followed by Bonferroni’s multiple comparisons test.

## Discussion

Despite the paucity of Treg in the retina, in certain conditions, such as retinal neovascularization or uveitis, systemic Treg are actively recruited to the retina, where they influence microglial activation and attenuate disease severity(McPherson et al. 2012; Deliyanti et al. 2017). Aging is characterized by a low-grade chronic inflammation, and a marked gliosis in different areas of the CNS including the brain. Recent work showed that local expansion of Treg in the brain through interleukin-2 overexpression reverses molecular and cognitive signatures of aging such as gliosis, inflammation, and cognitive decline(Lemaitre et al. 2023). Similar to what has already been described in the brain, disruption of retinal immune regulation is a key contributor to the development of age-related diseases(Chen et al. 2019). The role of Treg has been extensively studied in the context of autoimmune and inflammatory eye diseases(Lee & Foulsham 2022), but its role in retinal aging and homeostasis in the absence of overt pathology remains unknown. In part due to the low number of Treg found in healthy retina, Treg have not been previously investigated in the context of the non-pathological aging retina. Our study has addressed this question revealing a novel physiological role of Treg in maintaining retinal homeostasis and thereby limiting age-associated degenerative changes. We have shown how Treg, are essential in the maintenance of a healthy age-related para-inflammation, since its depletion led to exacerbated damage in RPE, phagocyte accumulation and neurodegeneration in the outermost layers of the neuroretina. Additionally, we have shown that once immunoregulation in the aging retina is lost, young Treg are more efficient at limiting inflammation and neurodegeneration than aged Treg, suggesting an intrinsic functional deficit in aged Treg in this setting. This study has identified Treg as key players in limiting age-associated retinal neurodegeneration, opening a new therapeutic avenue to preventing retinal degeneration in aged-related eye diseases. Future work that identifies the molecular mechanisms of Treg-mediated retinal neuroprotection could help to harness the anti-inflammatory and neuroprotective potential of Treg to treat age-associated retinal diseases.

## Figure Legends

**Figure EV1:**
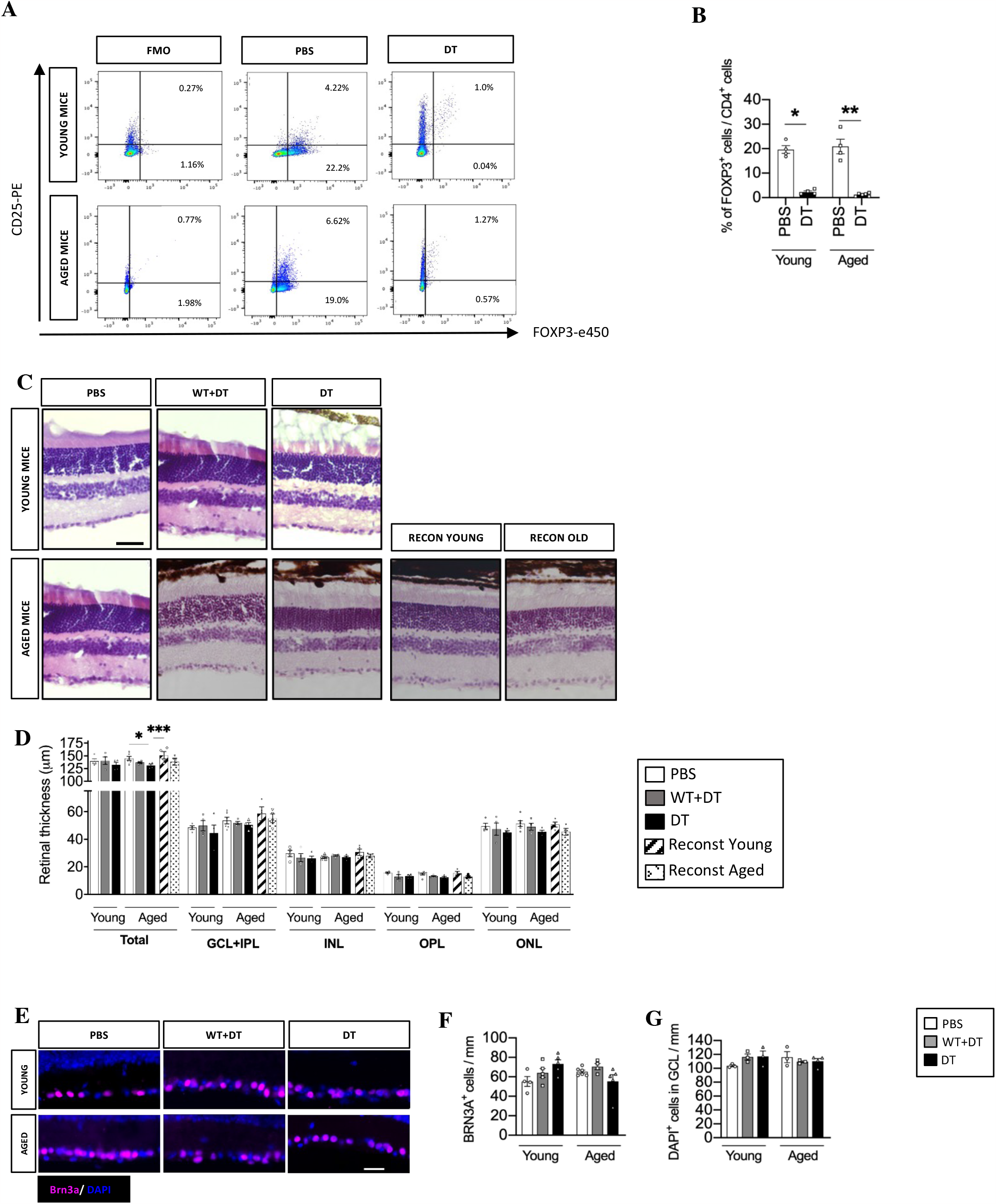
Validation of Treg depletion and additional neuroretinal characterization. **A, B** Determination of endogenous Treg in the lymph nodes in youth and aged Treg depleted animals. Flow cytometric plot (**A**) and analysis of endogenous Treg (**B**) in the lymph nodes identified by CD4, CD25 and Foxp3 staining to verify Treg depletion. **C, D** Hematoxylin and eosin analysis of retinal thickness. Representative images showing hematoxylin and eosin staining (**C**) of young and aged retinas in the different treatment groups (scale bar=50 μm). Quantification of the total retinal layer thickness as well as the thickness of the different cell layers (**D**). **E-G** Determination of RGC number in young and aged retinas. Representative image (E) (scale bar=25 μm) and quantification of retinal ganglion cells identified by Brn3a^+^ (**F**) and DAPI+ staining in GCL (**G**). Data information: A, B n=4-6 mice, C-G n=3-6. A-G data presented as mean±s.e.m. ^*^P<0.05; ^**^ P<0.01; ^***^P<0.005; ^****^p<0.001, U-Mann Whitney (B) and 1-way ANOVA followed by Bonferroni’s multiple comparisons test (D-G).

**Figure EV2:**
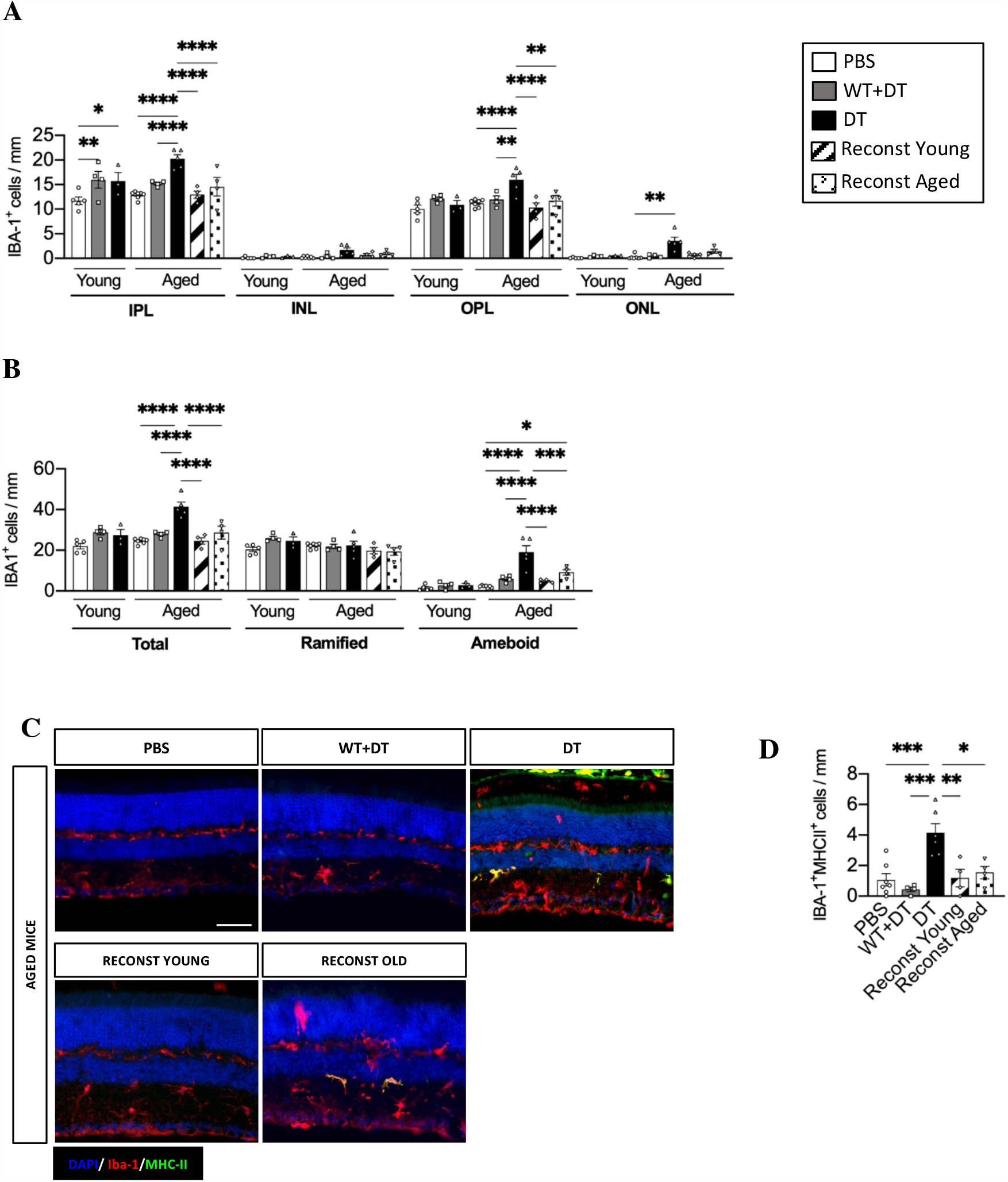
Analysis of microglia morphology, distribution, and phenotype. **A** Quantification of IBA-1^+^ microglia in the different layers of the neuroretina **B** Quantification of total, ramified, and ameboid microglia in the whole neuroretina of young and aged control and Treg depleted mice as well as aged-reconstituted mice. **C, D** Determination of IBA-1^+^MHCII^+^ cells in the neuroretina in aged mice. Representative image (**C**) (scale bar=50 μm) and quantification of IBA-1^+^MHCII^+^ microglia in the neuroretina of aged control, Treg depleted and reconstituted mice (**D**). Data information: A-D n=3-6 mice, data presented as mean±s.e.m. ^*^P<0.05; ^**^ P<0.01; ^***^P<0.005; ^****^p<0.001, 1-way ANOVA followed by Bonferroni’s multiple comparisons test.

## Methods

### Animals

All animals were housed and bred in a standard pathogen free experimental facility and exposed to a 12 h light/dark cycle with free access to food and water. All mice were bred in-house or purchased from Charles River Laboratories, UK. All procedures were conducted under the regulation of the UK Home Office Animals (Scientific Procedures) Act 1986. This study was approved by the Animal Welfare and Ethical Review body (AWERB) of Queen’s University Belfast and ethical review committee and all animal maintenance and experiments were done in according with the UK Home Office Regulation (Project Licenses 2789 and 2894). Foxp3-DTR mice were kindly provided by Prof. Alexander Rudensky (Memorial Sloan Kettering Institute, New York) and Dr. Rebecca Ingram (Queen’s University Belfast) and bred on a C57BL6/J background. Following the 3Rs principle and to reduce animal use, the eyes analyzed in this project were obtained from animals that underwent demyelination surgery exclusively in the spinal cord as described previously^9^. In brief, all mice underwent 30 min of isofluorane-based anaesthesia to allow spinal cord surgery, during which the eyes were covered with eye drops to prevent dehydration. Mice received a single injection of 1.2μl of 1% (w/v) L L-α-Lysophosphatidylcholine (Lysolecithin; Sigma-Aldrich) into the thoracic spinal cord under general anaesthesia. At day 18 post first DT injection (or day 14 post-surgery), mice were terminally anaesthetised with intraperitoneal (i.p.) pentobarbital injections and transcardially perfused with ice-cold phosphate buffered saline (PBS) followed by 4% paraformaldehyde (PFA) (Sigma-Aldrich). Eyes were removed and immersed in 4% PFA overnight at 4 °C. Then, eyes reserved for cryosectioning were cryoprotected with 30% sucrose in PBS for 72h, snap-frozen in OCT (Tissue-Tek), cryosectioned at 15 μm thickness and immunostained as indicated below. Eyes that were used for flatmount staining were fixed overnight and transferred to PBS.

### Treg depletion

Endogenous Treg were depleted from young (2-4m) and aged (18-23m) male and female Foxp3-DTR mice using diphtheria toxin (DT). Mice received daily i.p. injections of DT (0.04 μg/g of body weight; Sigma, Cat. No. D0564) for 3 consecutive days. Young (2-4m) and aged (16m) male and female C57BL6/J were used as controls for DT-associated side effects. To maintain endogenous Treg depletion throughout the course of the study, all mice received an i.p. DT injection (0.04μg/g of body weight) every fourth day up until day 18, when mice were sacrificed. Control animals received 200 μl of saline i.p. Depletion was confirmed at the endpoint by flow cytometric analysis of Foxp3 expression in blood, spleen, and lymph nodes.

### Natural Treg isolation and adoptive transfer

Young (2-4m) and aged (15-18m) female mice were culled by CO_2_ overdose. Lymph nodes and spleens were dissected and mashed into a single cell suspension using a 5ml syringe plunger. Single cell suspension was passed through a 70 μm strainer and then subjected to CD4 negative and CD25 positive immunomagnetic selection following manufacturer’s instructions. In brief, cells were incubated with CD4 negative selection kit (STEMCELL Technologies) for 15 min on ice and CD4^-^ cells were magnetically removed. Then, CD4^+^ cells were subjected to CD25^+^ cell isolation (STEMCELL Technologies). Cells were resuspended at 10^6^ cells per 200μl saline and injected i.p. on the day of the third DT injection (the day prior to spinal cord surgery).

### Flow cytometry

To confirm cell purity isolated CD4^+^CD25^+^ Treg were subjected to flow cytometry staining as described below. To confirm Treg depletion spleen and lymph nodes were mashed through a 70 μm strainer and then treated with red blood cell lysis buffer (STEMCELL Technologies) for 2 min at room temperature. Cells obtained from lymph nodes and spleens were then washed with PBS and centrifuged at 300 g for 5 min at 4°C. Cells were resuspended in 200 μL PBS and stained with a cell viability dye with eFluor 455-UV viability dye (1:2000; ThermoFisher Scientific) and cell surface stained with antibodies for CD4 (1:500; eBioscience, clone RM4.5) and CD25 (1:500; eBioscience, clone PC61.5) for 15 min at RT. Cells were washed with flow cytometry staining buffer (FCSB) (2% FCS in PBS) and centrifuged at 300 g for 5 min at 4 °C. Cells were then fixed with Fix & Perm A (ThermoFisher Scientific) for 10 min at RT. Fixative was washed off with FCSB and centrifuged at 300 g for 5 min at 4 °C. Then, cells were resuspended in 100 μL Fix & Perm B (ThermoFisher Scientific) with an anti-Foxp3 antibody (1:100; eBioscience, clone FJK-16S) overnight at 4 °C. Cells were then washed with FCSB and centrifuged at 300 g and 4 °C for 5 min. Final pellet was resuspended, data were acquired on a FACSCanto II and analyzed using FlowJo software version 9.0 (BD). To calculate cell numbers, singlets were identified by FSC-H versus FSC-A and viable cells gated on CD4 expression, and subsequently CD25^+^ and Foxp3^+^ and CD25^+^Foxp3^+^ cells.

### Immunostaining

#### Cryosection staining

Eye sections were dried for 30 min at RT and washed in PBS for 10 min. Antigen retrieval was performed using Citrate buffer pH 6.0 (Abcam, ab93678) at 95ºC for 5 minutes in a water bath and after cooling, and additional 5 minutes with 10% SDS. Slides were then blocked in 10% Donkey serum (Sigma Aldrich, D9663) in 0.2% Triton-X in PBS for 1h at room temperature. Primary antibodies (Table 1) were added and incubated overnight at 4ºC in 0.5% donkey serum and 0.2% triton-X followed by incubation with secondary antibodies (Table 1) for 1h at room temperature, in PBS.

#### Retinal Pigment Epithelium (RPE)/Choroid Flatmount staining

RPE/choroid flatmounts were dissected under a microdissection microscope (Nikon smz800, Nikon, Tokyo, Japan). The anterior segment of the eye (cornea, lens, iris and ciliary body) was removed, and the retina was carefully peeled off of the RPE/choroidal eyecup. RPE/choroid flatmounts tissues were washed in PBS followed by treatment with 2% Triton-X 100 for 2 h at RT. After washing, samples were blocked with 5% Donkey serum albumin in 0.2% Triton-X 100 for 1 hour at RT, and then incubated with the corresponding primary antibody and secondary antibodies shown in Table 1.

#### Image acquisition and analysis

The samples were cover-slipped with Vectashield (H-1000-10, Vector Labs, Burlingame, CA) and examined by Leica DMi8, DM550 epifluorescence microscope or confocal microscope (Leica TCS SP5 and SP8, Leica Microsystems Ltd., Wetzlar, Germany). Further image processing and analysis was performed using Fiji^25^ software and blindly manual counting.

### Statistical analysis

Statistical analysis was performed using Graph Pad Prism (GraphPad Software, Inc. version 9). First, normal distribution was assessed using Kolmogorov-Smirnov tests. When only two groups were compared, such as when checking Treg percentage changes upon Treg depletion U-Mann Whitney was used. For comparisons involving more than two groups, 1-way ANOVA analysis was performed followed by Bonferroni’s multiple comparisons test. When the data was described as percentage, an arcsin conversion was performed to analyze the data using parametric tests for normally distributed data. For all statistical tests, differences were considered significant with p values below 0.05.

## Author Contributions

Experiments were designed by M.L.S, A.G.F, A.S and D.C.F. Experiments were performed by M.L.S, A.G.F, and R.P. Experiments were analyzed by M.L.S, A.G.F and C.E.M. R.P and Y.D provided advice on experimental design and data interpretation. Manuscript was written by M.L.S, A.G.F, A.S and D.C.F with contributions from all authors. A.G.F and A.S oversaw the study.

## Acknowledgements

We acknowledge extensive technical support from the staff of the animal facility and Advanced Imaging CTU of the Wellcome-Wolfson Institute for Experimental Medicine. We thank A. Rudensky (Memorial Sloan Kettering Cancer Centre) and Rebecca J Ingram (Queen’s University Belfast) for providing Foxp3-DTR mice. This work was supported by the Wellcome (110138/Z/15/Z to DCF), ECTRIMS postdoctoral fellowship (to AGF), Wellcome Trust ISSF fellowship through QUB (to AGF), Miguel Servet Fellowship from the Spanish Institute of Health Carlos III (CP21/00032 to AGF), The Leverhulme Trust (ECF-2014-390, to YD), the Maria Zambrano fellowship from Spanish Ministry of Science, Innovation and Universities, financed by European Union “NextGenerationEU” (Universitat Autònoma de Barcelona, to MLS), and H2020 RECOGNISED, European Commission and Fight for Sight UK Grant Agreement N° 847749 (to AS).

## Data availability and citation experiment

All original data is available from the corresponding authors upon request.

**Primary antibody table-Table 1:**
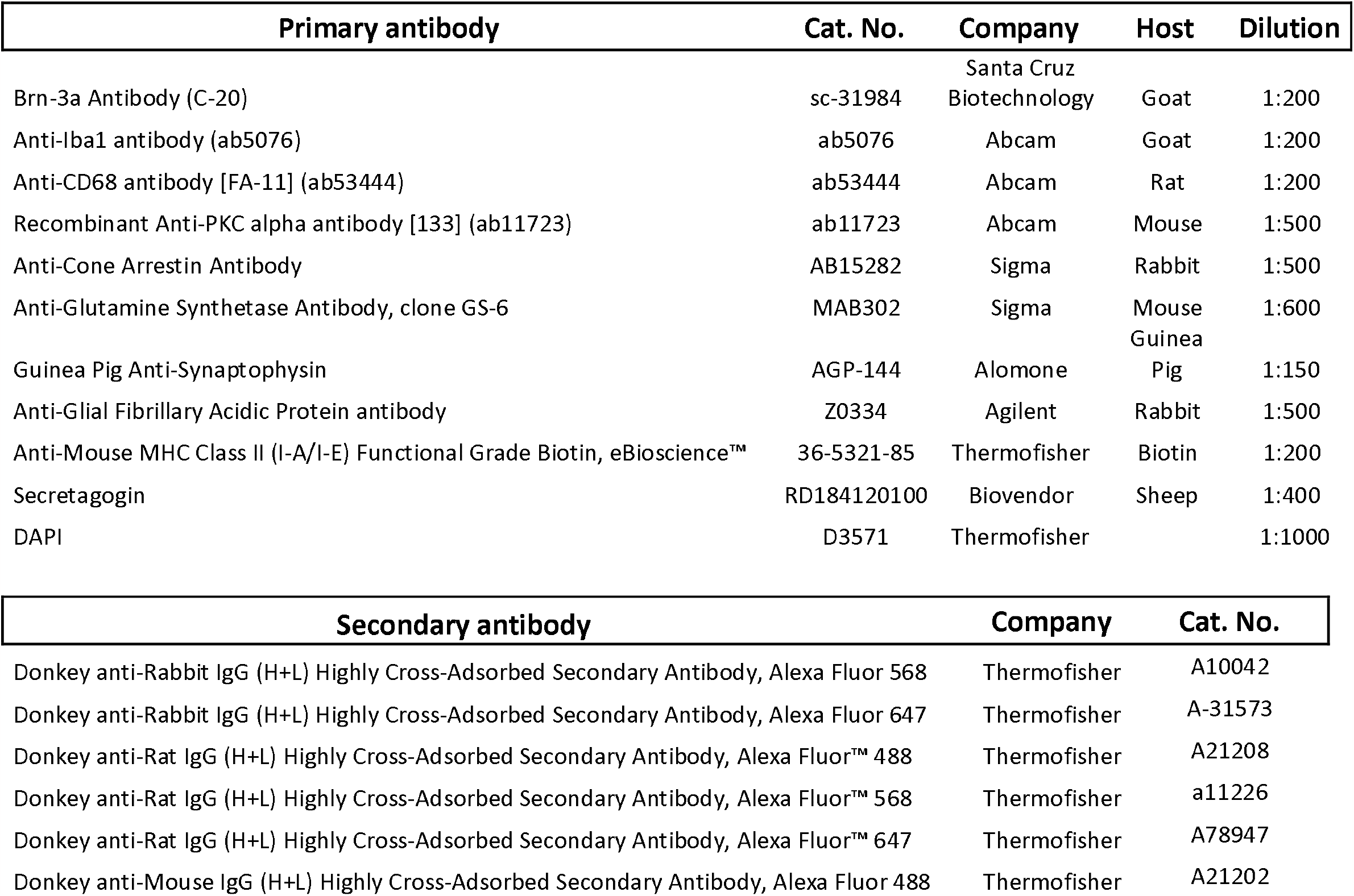
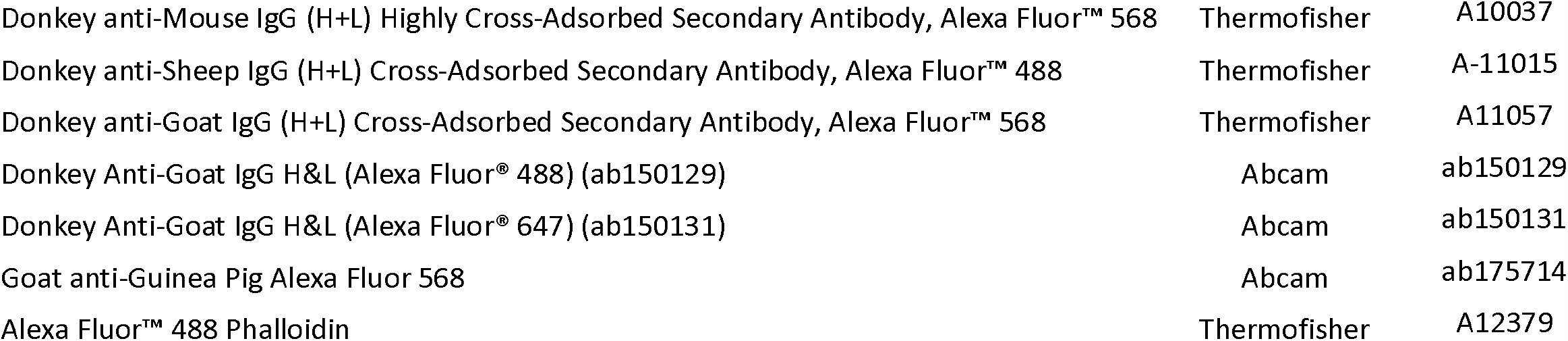

## References

Bringmann A, Iandiev I, Pannicke T, Wurm A, Hollborn M, Wiedemann P, Osborne NN & Reichenbach A(2009) Cellular signaling and factors involved in Müller cell gliosis: Neuroprotective and detrimental effects. Progress in retinal and eye research 28, 423–451. 10.1016/j.preteyeres.2009.07.001.

Butovsky O & Weiner HL(2018) Microglial signatures and their role in health and disease. Nature Reviews Neuroscience 19, 622–635. 10.1038/s41583-010-0057-5.

Chen M, Forrester JV & Xu H(2011) Dysregulation in Retinal Para-Inflammation and Age-Related Retinal Degeneration in CCL2 or CCR2 Deficient Mice. PLoS ONE 6, e22818. 10.1371/journal.pone.0022818.

Chen M, Luo C, Zhao J, Devarajan G & Xu H(2019) Immune regulation in the aging retina. Progress in retinal and eye research 69, 159–172. 10.1016/j.preteyeres.208.10.003.

Cuenca N, Fernández-Sánchez L, Campello L, Maneu V, Villa PD la, Lax P & Pinilla I(2014) Cellular responses following retinal injuries and therapeutic approaches for neurodegenerative diseases. Progress in retinal and eye research 43, 17–75. 10.1016/j.preteyeres.2014.07.001.

Deliyanti D, Talia DM, Zhu T, Maxwell MJ, Agrotis A, Jerome JR, Hargreaves EM, Gerondakis S, Hibbs ML, Mackay F & Wilkinson-Berka JL(2017) Foxp3+ Tregs are recruited to the retina to repair pathological angiogenesis. Nature Communications 8, 748. 10.1038/s41467-017-00751-w.

Dombrowski Y, O’Hagan T, Dittmer M, Penalva R, Mayoral SR, Bankhead P, Fleville S, Eleftheriadis G, Zhao C, Naughton M, Hassan R, Moffat J, Falconer J, Boyd A, Hamilton P, Allen IV, Kissenpfennig A, Moynagh PN, Evergren E, Perbal B, Williams AC, Ingram RJ, Chan JR, Franklin RJM & Fitzgerald DC(2017) Regulatory T cells promote myelin regeneration in the central nervous system. Nature Neuroscience 20, 674–680. 10.1038/nn.4528.

Elyahu Y, Hekselman I, Eizenberg-Magar I, Berner O, Strominger I, Schiller M, Mittal K, Nemirovsky A, Eremenko E, Vital A, Simonovsky E, Chalifa-Caspi V, Friedman N, Yeger-Lotem E & Monsonego A(2019) Aging promotes reorganization of the CD4 T cell landscape toward extreme regulatory and effector phenotypes. Science advances 5, eaaw8330. 10.1126/sciadv.aaw8330.

Fernández-Sánchez L, Lax P, Campello L, Pinilla I & Cuenca N(2015) Astrocytes and Müller Cell Alterations During Retinal Degeneration in a Transgenic Rat Model of Retinitis Pigmentosa. Frontiers Cell. Neurosci. 9, 484. 10.3389/fncel.2015.00484.

Frank MG, Barrientos RM, Biedenkapp JC, Rudy JW, Watkins LR & Maier SF(2006) mRNA upregulation of MHC II and pivotal pro-inflammatory genes in normal brain aging. Neurobiology of Aging 27, 717–722. 10.1016/j.neurobiolaging.

Garg SK, Delaney C, Toubai T, Ghosh A, Reddy P, Banerjee R & Yung R(2014) Aging is associated with increased regulatory T-cell function. Aging Cell 13, 441–448. 10.1111/acel.12191.

Hippert C, Graca AB, Barber AC, West EL, Smith AJ, Ali RR & Pearson RA(2015) Müller Glia Activation in Response to Inherited Retinal Degeneration Is Highly Varied and Disease-Specific. Plos One 10, e0120415. 10.1371/journal.pone.0120415.

Ito M, Komai K, Mise-Omata S, Iizuka-Koga M, Noguchi Y, Kondo T, Sakai R, Matsuo K, Nakayama T, Yoshie O, Nakatsukasa H, Chikuma S, Shichita T & Yoshimura A(2019) Brain regulatory T cells suppress astrogliosis and potentiate neurological recovery. Nature 565, 246–250. 10.1038/s41586-018-0824.

Kaur G & Singh NK(2021) The Role of Inflammation in Retinal Neurodegeneration and Degenerative Diseases. International Journal of Molecular Sciences 23, 386. 10.3390/ijms23010386.

Kim JM, Rasmussen JP & Rudensky AY(2007) Regulatory T cells prevent catastrophic autoimmunity throughout the lifespan of mice. Nature Immunology 8, 191–197. 10.1038/ni1428.

Lee AY & Foulsham W(2022) Regulatory T Cells: Therapeutic Opportunities in Uveitis. Frontiers Ophthalmology 2, 901144. 10.3389/fopht.2022.901144.

Lemaitre P, Tareen SH, Pasciuto E, Mascali L, Martirosyan A, Callaerts-Vegh Z, Poovathingal S, Dooley J, Holt MG, Yshii L & Liston A(2023) Molecular and cognitive signatures of ageing partially restored through synthetic delivery of IL2 to the brain. Embo Molecular Medicine, 15, e16805. 10.15252/emmm.202216805.

Madry C, Kyrargyri V, Arancibia-Cárcamo IL, Jolivet R, Kohsaka S, Bryan RM & Attwell D(2018) Microglial Ramification, Surveillance, and Interleukin-1β Release Are Regulated by the Two-Pore Domain K+ Channel THIK-1. Neuron 97, 299–312.e6. 10.1016/j.neuron.2017.12.002.

McPherson SW, Heuss ND & Gregerson DS(2012) Regulation of CD8+ T Cell Responses to Retinal Antigen by Local FoxP3+ Regulatory T Cells. Frontiers Immunology 3, 166. 10.3389/fimmu.2012.00166.

Muñoz-Rojas AR & Mathis D(2021) Tissue regulatory T cells: regulatory chameleons. Nature Review Immunology 21:597–611. 10.1038/s41577-021-00519-w.

Shi L, Sun Z, Su W, Xu F, Xie D, Zhang Q, Dai X, Iyer K, Hitchens TK, Foley LM, Li S, Stolz DB, Chen K, Ding Y, Thomson AW, Leak RK, Chen J & Hu X(2021) Treg cell-derived osteopontin promotes microglia-mediated white matter repair after ischemic stroke. Immunity 54, 1527–1542.e8. 10.1016/j.immuni.2021.04.022.

Tarau I-S, Berlin A, Curcio CA & Ach T(2019) The Cytoskeleton of the Retinal Pigment Epithelium: from Normal Aging to Age-Related Macular Degeneration. International Journal of Molecular Sciences 20, 3578. 10.3390/ijms20143578.

Xu H, Chen M & Forrester JV(2009) Para-inflammation in the aging retina. Progress in retinal and eye research 28, 348–368. 10.1016/j.preteyeres.2009.06.001.

Yshii L, Pasciuto E, Bielefeld P, Mascali L, Lemaitre P, Marino M, Dooley J, Kouser L, Verschoren S, Lagou V, Kemps H, Gervois P, Boer A de, Burton OT, Wahis J, Verhaert J, Tareen SHK, Roca CP, Singh K, Whyte CE, Kerstens A, Callaerts-Vegh Z, Poovathingal S, Prezzemolo T, Wierda K, Dashwood A, Xie J, Wonterghem EV, Creemers E, Aloulou M, Gsell W, Abiega O, Munck S, Vandenbroucke RE, Bronckaers A, Lemmens R, Strooper BD, Bosch LVD, Himmelreich U, Fitzsimons CP, Holt MG & Liston A(2022) Astrocyte-targeted gene delivery of interleukin 2 specifically increases brain-resident regulatory T cell numbers and protects against pathological neuroinflammation. Nature Immunology 23, 878–891. 10.1038/s41590-022-01208-z.

